# Loss of physiological stretch elicits an inflammatory, plaque-like transcriptional state in cultured human aortic smooth muscle cells

**DOI:** 10.1101/2025.10.29.685276

**Authors:** Lise Filt Jensen, Anton Markov, Laura Alonso-Herranz, Diana Sharysh, Emil Aagaard Thomsen, Jacob Giehm Mikkelsen, Jacob Fog Bentzon, Julián Albarrán-Juárez

## Abstract

**Background and aims:** Vascular smooth muscle cells (SMCs) cultured under standard static conditions adopt a modulated phenotype that resembles the SMC states found in atherosclerotic plaques, but the mechanical determinants of this drift are not well defined. We investigated how mechanical loading shapes SMC phenotype and inflammatory signaling.

**Methods:** Human aortic SMCs were maintained under static conditions, physiological cyclic stretch (10% elongation), or pathological stretch (15% elongation) and analyzed by bulk and single-cell RNA sequencing. Mechanistic experiments included siRNA-mediated knockdown of *IKBKB*, p65 immunofluorescence, and regulon inference from single-cell transcriptomes.

**Results:** Stretch regulated cell cycle, contractile, and inflammatory gene programs in an intensity-dependent manner: 10% stretch suppressed basal and TNF-induced inflammatory gene expression, whereas 15% stretch did not. The anti-inflammatory effect of physiological stretch required IKBKB, yet the proportion of cells with nuclear p65 was unchanged, indicating that stretch constrains NF-κB output downstream of p65 nuclear entry rather than by blocking translocation. Single-cell RNA sequencing resolved nine states for SMCs in culture whose transcriptomes overlapped substantially with the modulated mesenchymal populations of human coronary and carotid plaques, and physiological stretch attenuated pro-inflammatory gene expression across the major clusters. Regulatory network analysis identified inflammatory transcription factors (RELB, CEBPB/D, IRF1/2, STAT2) as less active under physiological stretch, whereas mechano-lineage regulators (MEF2C, TEAD1, SMAD6) were selectively induced, providing candidate mediators of the effect.

**Conclusions:** Static culture represents a disease-like SMC baseline that physiological stretch attenuates, in part by constraining NF-κB transcriptional output at a step downstream of p65 nuclear entry.

**Graphical abstract:** Physiological stretch (10%) maintains a low-inflammatory, healthy-like state in human aortic smooth muscle cells, whereas its absence (static culture) or excess (15% stretch) favors an inflammatory, plaque-like state. NF-κB-p65 enters the nucleus under all conditions; physiological stretch constrains its inflammatory output downstream of nuclear entry.

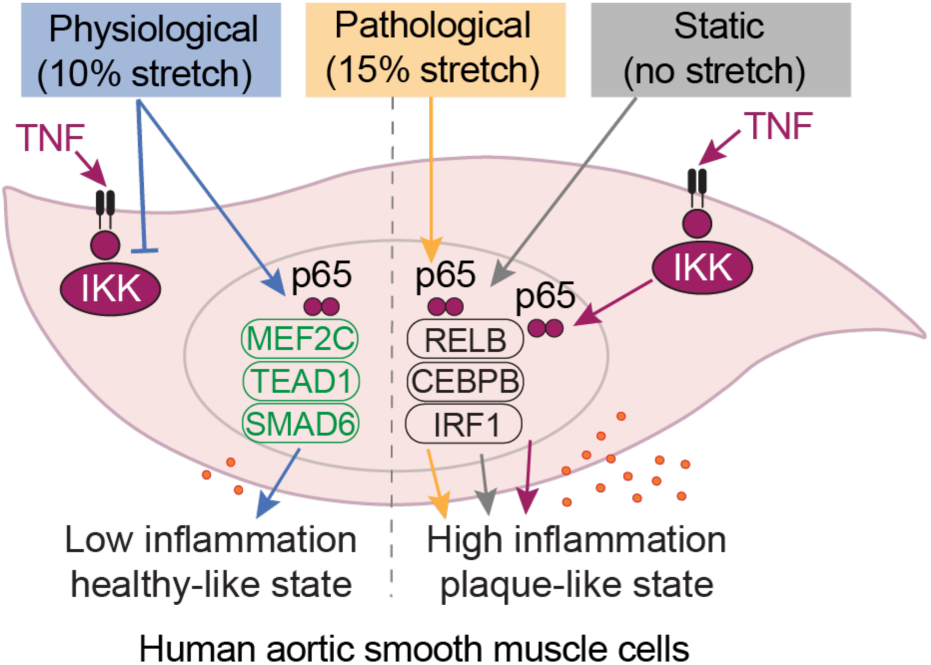

## 1. Introduction

Cardiovascular disease is the leading cause of mortality worldwide, and atherosclerosis, a chronic inflammatory condition of the arteries, is responsible for over 80% of these deaths^1^. During atherosclerosis, vascular smooth muscle cells (SMCs) undergo phenotypic modulation from a contractile to heterogeneous states expressing genes characteristic of other cell types, accompanied by decreased expression of contractile proteins and increased production of extracellular matrix (ECM) components^2–5^. Cultured SMCs adopt strikingly similar modulated states, and this overlap has long been attributed to the loss of physiological cues, in particular the cyclic mechanical forces experienced *in vivo*, which support the differentiated SMC phenotype^6–9^. In healthy elastic arteries, the vessel wall experiences approximately 10% cyclic stretch with each cardiac cycle^9–13^. During aging and atherosclerosis, extensive ECM remodeling leads to vessel stiffening and altered mechanical stresses^14,15^, with tensile forces concentrating in thin fibrous caps and plaque shoulders, regions prone to rupture^16–18^. This remodeling, characterized by increased deposition of interstitial collagens, fibronectin, osteopontin, and proteoglycans, further promotes SMC phenotypic modulation^19,20^. Inflammatory signaling, particularly through NF-κB, is a central driver of SMC modulation in atherosclerosis^21,22^, yet whether mechanical loading directly regulates this pathway in human SMCs remains unclear^23^. Here we combine bulk (RNA-seq) and single-cell RNA sequencing (scRNA-seq) with targeted perturbation to test whether mechanical stretch influences the modulated state that cultured SMCs adopt by default, and to define the transcription factors and associated programs through which it acts.

## 2. Materials and methods

### 2.1 Smooth muscle cell culture

Human aortic SMCs (ATCC, PCS-100-012, male donor, 26 years old) were authenticated by ATCC and tested negative for mycoplasma contamination prior to experiments. Cells were cultured in smooth muscle cell growth medium (SMC-GM) containing 5% fetal bovine serum (FBS), bFGF (2 ng/mL), EGF (0.5 ng/mL), insulin (5 µg/mL), gentamicin (50 µg/mL), and amphotericin B (50 ng/mL) (Provitro, 211-0601) at 37 °C and 5% CO_2_. Only cells in passages 4-6 were used for the experiments. All experiments were performed in growth medium (SMC-GM) as stated above.

### 2.2 Collagen coating

For plastic labware, plates were coated with collagen type I (COL1) from rat tail (Corning, 354236) diluted in 0.1% acetic acid at a concentration of 20 µg/mL for 20 min at room temperature (RT) and washed once with Phosphate-Buffered Saline (PBS). For stretch, precoated COL1 BioFlex plates were used (Flexcell, BF-3001C).

### 2.3 Mechanical stretch

SMCs were seeded onto BioFlex six-well plates (2#x00D7;10^5^ cells per well) and cultured to ∼80-90% confluency. Immediately before loading, the medium was replaced with fresh SMC-GM in all conditions, and stretch was then applied. Cyclic equibiaxial stretch was applied using the Flexcell tension system^24,25^ (FX-5000T, Flexcell International Corporation, Burlington, NC, USA) at 10% or 15% elongation, delivered as a sinusoidal waveform at 1 Hz for 6 h; control cells were held under static conditions.

### 2.4 Immunostaining

SMCs on BioFlex plates were washed with pre-warmed PBS and fixed with pre-warmed 4% paraformaldehyde (PFA) for 15 min at room temperature (RT). SMCs were washed three times with PBS and permeabilized with 0.1% Triton X-100 for 10 min at RT. Nonspecific binding was blocked with 3% bovine serum albumin (BSA) in PBS for 1 h at RT. For staining, the silicon membranes were cut out before continuing with antibody incubations in humidified chambers. Primary antibodies were diluted in 0.3% Triton X-100 and 1% BSA in PBS and applied as follows: mouse monoclonal anti-α-tubulin (1:2000, Sigma-Aldrich, clone B512, T5168, RRID:AB_477579) for 1 h at 37 °C; phalloidin conjugated to Alexa Fluor 568 (1:1000, Invitrogen, A12380) for 30 min at RT; rabbit monoclonal anti-MKI67 (SP6) (1:1000, Abcam, ab16667, RRID:AB_302459) overnight (ON) at 4 °C; rabbit monoclonal anti-NF-κB p65 (D14E12) (1:1000, Cell Signaling Technology, 8242, RRID:AB10859369) ON at 4 °C. Secondary antibodies, Alexa Fluor 488 donkey-anti-mouse IgG (H+L) (1:800, Invitrogen A21202) or Alexa Fluor 488 donkey-anti-rabbit IgG (H+L) (1:800, Invitrogen A21206), were incubated for 1 h, at RT. DNA was stained using DAPI (2.5 ng/mL; Invitrogen, D1306) in PBS for 10 min at RT. BioFlex membranes were mounted using Fluoromount^TM^ aqueous mounting medium (Sigma-Aldrich, F4680). Representative images (3-6 per replicate) were acquired using an inverted fluorescent microscope (Nikon eclipse Ti2) with a Nikon CFI S plan Fluor ELWD ADM 20x objective, NA=0.45 and analysis was performed in ImageJ^26^. MKI67 was quantified as the percentage of MKI67-positive cells relative to the total number of cells. NF-κB p65 nuclear localization was assessed by calculating the nuclear-to-cytoplasmic mean fluorescence intensity (MFI) ratio for each cell. Cells with a ratio >1 were classified as exhibiting nuclear localization, whereas cells with a ratio <1 were classified as exhibiting cytoplasmic localization. For each condition, the percentage of cells exhibiting nuclear localization was calculated relative to the total number of quantified cells.

### 2.5 RNA isolation and cDNA synthesis

RNA was isolated using the NucleoSpin RNA Plus Kit (Macherey-Nagel, 790984). RNA quantity and quality were assessed using a Nanodrop One spectrophotometer (ThermoFisher Scientific). cDNA was synthesized from RNA using the RevertAid First Strand cDNA Synthesis kit (ThermoFisher Scientific, K1622), according to the manufacturer’s instructions.

### 2.6 Bulk RNA sequencing and bioinformatics analysis

For RNA sequencing three independent samples per condition were used. The bulk RNA (transcriptome) was sequenced by BGI Genomics (Copenhagen, Denmark). All RNA samples passed quality control (Agilent 4200 TapeStation electrophoresis system). Non-stranded cDNA libraries were prepared with polyA-selected mRNA, and 100 bp paired-end sequencing was conducted with at least 20 million paired reads per sample using the DNBSEQ-G400 platform. RNA sequencing data were processed with the Nextflow-based pipeline “nf-core/rnaseq” (version 3.9). Sequenced reads were aligned to the GRCh38.p13 human reference genome using the STAR aligner (version 2.7.10a), and gene-level quantification was done with Salmon (version 1.5.2). The reference genome sequence and gene annotation were downloaded from GENCODE (release 41). Further analysis was performed in R (version 4.4.2, https://www.r-project.org) with Bioconductor (version 3.20) packages^27^. Gene counts were imported using tximport^28^. Gene filtering was performed, and only genes with ≥10 counts in at least 3 samples were analyzed. DESeq2^29^ was used for data normalization, principal component analysis, and estimation of differential gene expression between compared groups. Results with false discovery rate adjusted p-values (padj) <0.05 and log2 fold-change greater than 0.5 or less than −0.5 were considered differentially expressed and statistically significant.

### 2.7 Real-time qPCR

Relative gene expression was measured by real-time quantitative PCR (RT-qPCR). Intron-spanning primers rendering 50-150 bp amplicons were designed **(Supplementary Table 1)**. The Maxima SYBR Green qPCR master mix kit, including ROX (0.18 µM, Thermo Fisher Scientific, K0252), was mixed with forward and reverse primers (300 nM each) and cDNA (12.5 ng) in a final volume of 14 µL. Samples were run in duplicate. The RT-qPCR program was run on a Bio-Rad CFX Opus instrument as follows: 1 cycle of 95 °C for 10 min; 40 cycles of 95 °C for 30 sec, 60 °C for 1 min, and 72 °C for 1 min; followed by a melt curve step (95 °C for 1 min, 60 °C for 30 sec, and 95 °C for 30 sec). Relative mRNA expression was calculated using the ΔΔCt method with *HPRT1* as the reference gene and static untreated samples as the control samples.

### 2.8 Pathway enrichment analysis

Enrichment analysis of the RNA-seq data was performed using the WEB-based Gene SeT AnaLysis Toolkit (WebGestalt)^30^ with the Kyoto Encyclopedia of Genes and Genomes (KEGG) database^31^. The submitted gene lists are shown in **Supplementary Table 2-3 and 6**, for bulk RNA-seq and scRNA-seq analyses, respectively. Only genes with padj <0.05 and log2 fold-change greater than 0.5 or less than −0.5 were considered differentially expressed, and over-representation analysis was used.

### 2.9 Proliferation assay

To label proliferating cells, SMCs were incubated with 5-ethynyl-2’-deoxyuridine (EdU) (Sigma-Aldrich BCK-EDU488) for 6 h. After that, the cells were fixed and stained according to the manufacturer’s instructions, and all nuclei were stained with DAPI (2.5 ng/mL) in PBS. BioFlex silicon membranes were cut out and mounted using Fluoromount aqueous mounting medium. Representative images (4-8 per independent replicate) were acquired using an inverted fluorescent microscope (Nikon Eclipse Ti2) with a Nikon CFI S Plan Fluor ELWD ADM 20x objective, NA=0.45. The percentage of proliferating cells (%EdU+ cells) was calculated as the number of EdU-positive cells divided by the total number of cells. Each data point represents the average %EdU+ cells per replicate.

### 2.10 Collagen I gel contraction assay

Human aortic SMCs were cultured on COL1-coated BioFlex plates and exposed to either static conditions or 10% stretch for 6 h. A gel contraction assay was then performed as described^32^. Stretched SMCs were harvested from BioFlex plates and resuspended at 5#x00D7;10^5^ cells/mL in SMC-GM. COL1 gels were prepared by mixing rat tail collagen-I (Corning, 354236) with 0.1% acetic acid to a final concentration of 3 mg/mL and kept on ice. SMCs were mixed with the collagen-I solution at a 1:1 ratio, and the pH was neutralized by addition of 1 M NaOH (11.67 µL per mL solution). 0.5 mL of the cell–collagen mixture was added per well of a low-adherence 24-well plate (Falcon, 10396881), corresponding to 1.25#x00D7;10^5^ cells per gel. Collagen gels without SMCs were included as negative controls. The gels were polymerized at 37°C and 5% CO_2_ for 30 min, after which 0.5 mL of medium with or without transforming growth factor β1 (TGFβ1, 20 ng/mL; PeproTech, 100-21) was added. Gels were detached from the well edges using a pipette tip and gently swirled to allow free-floating contraction. Gel areas were recorded at 24 h and 48 h using a stereomicroscope (Kern Optics, OZM903) coupled to a camera (Kern Optics, OCD832) and quantified in ImageJ^26^. Gel contraction was reported as the relative gel area normalized to the area of the negative control.

### 2.11 TNF treatment

Human recombinant tumor necrosis factor (TNF) protein (Abcam, ab259410) was diluted in SMC-GM at a final concentration of 10 ng/mL. SMCs were treated with TNF for 24 h, starting at time 0; during the final 6 h, cells were subjected to static conditions, 10% stretch, or 15% stretch.

### 2.12 Small interference RNA (siRNA) knockdown

*IKBKB* knockdown (KD) was performed using a pool of two different siRNAs targeting human *IKBKB*: Hs_IKBKB_8 (S102626456) and Hs_IKBKB_14 (S105460693) (20 nM final concentration, QIAGEN). Cells were transfected with Lipofectamine RNAiMAX (Invitrogen, 13778150) on two consecutive days following the manufacturer’s guidelines. Treatments were started 24 h after the final transfection. A scramble siRNA (20 nM final concentration, AllStars Negative S103650318, QIAGEN) was used as a control.

### 2.13 NF-κB-EGFP reporter assay

An NF-κB reporter cassette in a lentiviral vector was kindly donated by Martin Schwartz and Brian Coon (Yale University, USA). This cassette consisted of three NF-κB response elements (GGGAATTTCC) controlling the activation of a minimal human cytomegalovirus (CMV_min_) promoter to drive the expression of a destabilized enhanced green fluorescent protein gene (d2EGFP). Additionally, the cassette contained a constitutively expressed fluorescent protein (DsRed) as an internal control. To enable antibiotic selection, the DsRed gene was removed by Gibson assembly and replaced with a blasticidin (Blast) resistance gene. Lentiviral production was performed as previously described^33^. SMCs were transduced at a multiplicity of infection (MOI) of 0.2 by adding the viral supernatant to SMC-GM supplemented with polybrene (8 µg/mL). To assess NF-κB activity, SMCs carrying the modified reporter cassette were stimulated with TNF (10 ng/mL) for 24 h or left untreated, washed twice with PBS, detached using 0.25% trypsin-EDTA (Gibco), and resuspended in PBS. Cells were analyzed on a NovoCyte 2100YB flow cytometer with two lasers (488 nm and 561 nm). To detect d2EGFP expression, the 488 nm laser and 530/30 filter were used (Agilent, Santa Clara, CA). The data were analyzed in Novo Express v1.5.6.

### 2.14 Single-cell RNA sequencing analysis

To investigate the heterogeneity of human aortic SMCs in culture, we performed scRNA-seq. Cells were cultured on BioFlex plates coated with collagen-I (COL1) under three conditions: static, 10% stretch, and TNF-stimulation (10 ng/mL). SMCs for all three conditions were seeded on the same day, and the following day, TNF stimulation for 24 h was initiated. To align endpoints, 10% stretch was applied during the final 6 h. Thus, the 24 h static condition serves as a control for all conditions. Stretch (6 h) and TNF (24 h) differed in duration by design, with the static condition as the common reference. Three samples from each condition were pooled, harvested, resuspended as single cell suspensions in PBS, and captured using the BD Rhapsody™ microwell-based cell capture platform. We followed the BD Rhapsody HT Single-Cell Analysis System (23-24257(01)) and Single-Cell Capture and cDNA Synthesis (23-24252(01)) protocols. Briefly, for each condition, 40,000 cells were loaded per lane, resulting in capture of ∼28,000 cells with ∼7–8% multiplet rate. Beads containing cell barcodes, unique molecular identifiers (UMIs), and oligo-dT sequences were loaded to each lane and used to capture polyadenylated mRNA after cell lysis directly in the microwells, followed by cDNA synthesis. Library preparation of ∼14,000 cells (half of each condition) was performed according to the BD Rhapsody™ Whole Transcriptome Analysis (WTA) protocol (23-24117(02)), and quality was confirmed using an Agilent Bioanalyzer. As forward primer for the index PCR, we used the TruSeq D501 primer, while the Nextera N709, N710, and N707 were used as reverse primers for the static, TNF, and 10% stretch libraries, respectively. Indexed libraries were pooled in equimolar amounts and sequenced on Illumina NovaSeq X Plus (2 × 150 bp, 10B flow cell, one lane) at OMIICS (Aarhus, Denmark), yielding ∼2–2.5 billion paired-end reads, corresponding to an expected read-pair per cell of ∼50,000. RNA sequencing data were processed and analyzed on the GenomeDK cluster. Raw data quality control and generation of a gene expression matrix were carried out following the BD Rhapsody™ Sequence Analysis Pipeline (23-24580(01)) protocol, with local pipeline software version 2.3 setup and basic cell detection algorithm. Further analysis was performed in R (version 4.4.2). and Seurat package (version 5.2.1)^34^. Cells having outlying decimal logarithms of transcripts and genes per cell (beyond five median absolute deviations from the median) were discarded. We also filtered out the cells having more than 15% of mitochondrial RNA content or mito-ribo ratio (the proportion of mitochondrial RNA in the total sum of mitochondrial and ribosomal read counts) above 0.75^35,36^. Putative doublets were detected using scDblFinder^37^ with 7% expected multiplet rate. Data from different conditions were normalized using the “LogNormalize” method (for each cell, gene counts are divided by the total counts, multiplied by 10,000, followed by natural log-transformation), and integrated in Seurat using a reciprocal principal component analysis (RPCA) approach with 15 principal components based on the expression matrix of 2000 most variable genes^38^. Cell clusters were identified using a shared nearest neighbor modularity optimization-based algorithm at varying resolutions (0.1 to 0.4). At a clustering resolution of 0.3, nine cell clusters were identified and retained for downstream analysis, as they corresponded to proliferating cells in various cell-cycle phases and to distinct quiescent populations. The Wilcoxon rank-sum test was used for both identification of marker genes and comparison between conditions. Cell cluster annotation was manually curated based on expressed marker genes.

### 2.15 Single-cell regulatory network inference and clustering (SCENIC)

Gene regulatory networks in human aortic smooth muscle cells subjected to static culture, 10% stretch, or TNF stimulation as previously mentioned were inferred from the integrated scRNA-seq dataset using the SCENIC workflow^39^. Raw counts were extracted from the integrated Seurat object and filtered to retain genes detected in at least five cells, applying default thresholds in SCENIC for minimum total UMI count (3 x 0.01 x total cell number) and minimum detection frequency (≥1% of cells); only genes present in the human RcisTarget^39^ motif databases were retained. Co-expression modules linking transcription factors to candidate target genes were inferred from the log2-transformed, filtered expression matrix using GENIE3. These modules were pruned to direct regulatory targets by cis-regulatory motif enrichment analysis in RcisTarget, yielding the final regulons. Per-cell regulon activity was scored on the transformed expression matrix using AUCell^39^, generating a continuous area-under-the-curve (AUC) matrix. Differential regulon activity was assessed between experimental groups (static, stretch, TNF) using global (one-vs-rest) comparison. Continuous AUC scores were compared using two-sided Wilcoxon rank-sum tests, restricted to regulons active (AUC > 0) in at least 10% of cells in either of the two compared groups; results are reported as mean AUC, log2 fold-change, and Benjamini-Hochberg (BH)-adjusted p-values. All analyses were performed in R (version 4.4.3).

### 2.16 Integration of scRNA-seq studies from public datasets

To map cultured SMCs to plaque SMCs identified in atherosclerotic lesions in coronary and carotid arteries, we collected scRNA-seq data publicly available^40–42^. External data preparation and processing steps are described elsewhere^43^. Then datasets were integrated using RPCA method with 30 principal components and preserving the original structure and annotation of cell clusters.

### 2.17 Statistical analysis

The statistical analysis was performed in GraphPad Prism, version 8.0.2 (GraphPad Software, Boston, Massachusetts USA). Samples were processed in parallel. Unless stated otherwise, all experiments were performed with ≥3 independent replicates. Sample size was based on previous studies and reproducibility considerations. Data are shown as the mean ± standard error of the mean (S.E.M.). Comparison between two groups was performed by a two-sided unpaired t-test if the data passed the Shapiro-Wilk normality test. Comparison of more than two groups, was done using one-way analysis of variance (ANOVA) followed by Tukey’s *post hoc* test. If the data did not pass the normality test, the non-parametric Kruskal-Wallis test was performed, followed by Dunn’s *post hoc* test. An adjusted p-value <0.05 was considered statistically significant, unless stated otherwise.

## 3. Results

### 3.1 Mechanical stretch intensity elicits distinct transcriptomic responses in human SMCs

To investigate how mechanical stretch alters the phenotype of human aortic SMCs, cells were cultured on COL1-coated soft silicone membranes and exposed to static conditions, 10% (physiological) or 15% (pathological) elongation for 6 hours prior to bulk RNA-seq **(Figure 1A, Supplementary Tables 2-3)**. These stretch levels were selected as previously described^43–46^. No major morphological or cytoskeletal differences were observed between conditions at this time **(Figure 1B)**, but principal component analysis revealed marked transcriptomic shifts **(Figure 1C)**. Differential gene expression analysis comparing 10% stretch with static condition **(Figure 1D)** and 15% with 10% stretch **(Figure 1E)** identified inflammatory signaling and cell cycle as the most significant enriched pathways. Relative to static culture, physiological stretch downregulated inflammatory marker genes, including adhesion molecules (*ICAM1, VCAM1*), chemokines (*CCL2, CCL8*), the signaling mediator *IRAK2*, and transcription factors (*CEBPD, RELB*); these genes were expressed at higher levels under pathological than physiological stretch. Cell cycle genes (e.g., *MKI67, CCNB1*) showed the inverse pattern, being upregulated by physiological stretch relative to static culture and expressed at lower levels under pathological stretch than 10%. However, only a few contractile markers (*CNN1, ACTG2*) were regulated under physiological stretch. Stretch intensity-dependent regulation was confirmed by RT-qPCR on a subset of genes (**Figure 1F-G**), supporting the RNA-seq findings. Together, these results indicate that stretch regulates inflammatory and cell cycle gene programs in an intensity-dependent manner, with physiological stretch suppressing inflammatory signaling and increasing cell cycle gene expression relative to static culture, an effect not sustained at pathological stretch.

**Figure 1:**
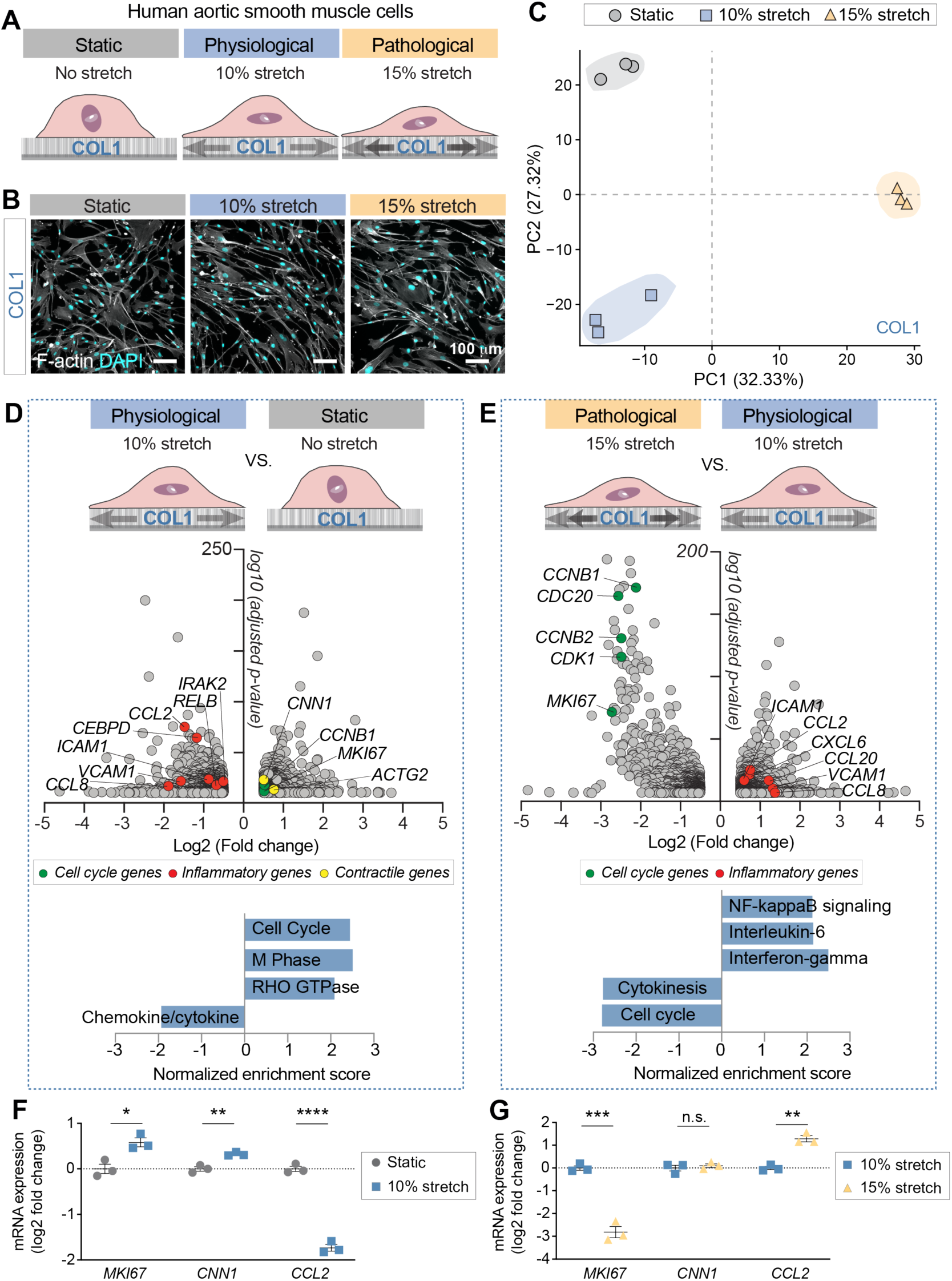
Mechanical stretch intensity regulates the transcriptome of SMCs. **A.** Schematic of the experimental design. SMCs were cultured on collagen-I (COL1)-coated surfaces followed by exposure to different mechanical stretch conditions (static, 10, or 15% elongation for 6 h, 1 Hz) and RNA-seq (*n*=3 samples per condition). **B.** Morphology and cytoskeleton (F-actin, gray color) of SMCs upon exposure to different stretch conditions. Nuclei are stained with DAPI (cyan). Scale bars: 100 μm. **C.** Principal component analysis (PCA) of transcriptome profiles of SMCs exposed to different stretch conditions (based on 37,483 genes). **D-E.** Volcano plots indicate differentially expressed genes (DEGs) with a false discovery rate adjusted p-values (padj) <0.05 and log2 fold-change greater than 0.5 or less than −0.5. Enrichment analysis of the RNA-seq data was performed using the WEB-based Gene SeT AnaLysis Toolkit (WebGestalt) with the Kyoto Encyclopedia of Genes and Genomes (KEGG) database. **D**. Illustrates the analysis comparing 10% stretch and static conditions, and **E.** 15% and 10% stretch conditions, accordingly. **F-G**. Relative mRNA expression of marker genes in SMCs subjected to different mechanical stretch conditions, determined by RT-qPCR and presented as log2 fold-change (*n*=3 samples per condition). The groups in F and G were compared by unpaired t-test. P-value: *p<0.05; **p<0.01; ***p<0.001; ****p<0.0001.

### 3.2 Physiological stretch suppresses NF-κB-dependent inflammatory signaling in SMCs

The most striking finding from the RNA-seq analysis was that physiological stretch downregulates multiple inflammation-responsive genes, several of which are primarily regulated by the NF-κB pathway. We therefore tested whether physiological stretch could also attenuate the effect of TNF, a classical NF-κB-mediated cytokine. TNF stimulation markedly increased *CCL2* and *ICAM1* expression in static SMCs, whereas this response was significantly blunted in cells exposed to 10% stretch **(Figure 2A).** In contrast, 15% stretch did not alter TNF-induced gene expression **(Figure 2B)**, reinforcing that this anti-inflammatory effect is tightly regulated with stretch intensity. To assess the role of NF-κB signaling, we combined TNF and physiological stretch with small interfering RNA (siRNA)-mediated knockdown of *IKBKB*, a key component of the canonical NF-κB pathway^47^. Physiological stretch alone reduced *IKBKB* expression, suggesting a potential mechanism by which physiological stretch may regulate inflammatory signaling **(Figure 2C)**. Moreover, 10% stretch did not further decrease *CCL2* expression in *IKBKB*-deficient cells, indicating that suppression of *CCL2* by physiological stretch depends on NF-κB signaling.

**Figure 2:**
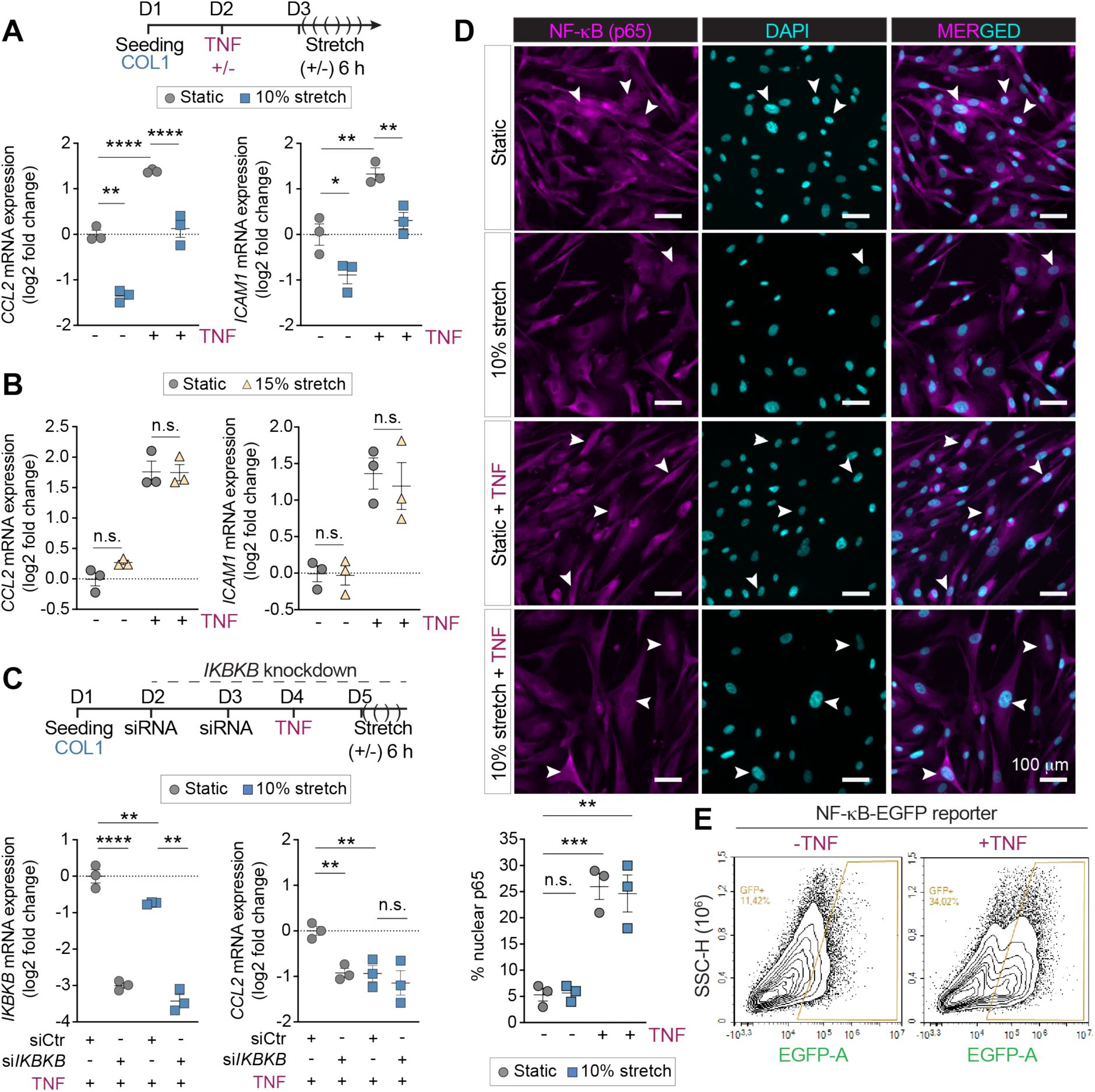
Physiological stretch attenuates inflammatory signaling in SMCs. **A-B.** Relative mRNA expression (log2 fold-change) measured by RT-qPCR of inflammatory genes (*CCL2, ICAM1)* in SMCs. Cells were either untreated or stimulated with TNF (10 ng/mL, 24 h), and subjected to static or 10% stretch **(A)** or 15% stretch **(B)**, respectively (*n*=3 independent replicates per condition). **C.** SMCs were transfected with negative control siRNAs (siCtr) or against *IKBKB* (si*IKBKB*), for two consecutive days. One day later, cells were treated with TNF for 24 h and then exposed to 10% stretch or left under static conditions. Shown is the relative mRNA expression measured by RT-qPCR of *IKBKB* and *CCL2* of three independent replicates. **D.** Representative images of p65 immunostaining of SMCs untreated or TNF-stimulated for 24 h and subjected to 10% stretch (6 h) or static conditions. The graph below shows the percentage of cells positive for nuclear p65 relative to the total number of cells (*n*=3 independent replicates per group). Scale bars: 100 µm. **E.** Representative plot of SMCs carrying the NF-κB-EGFP reporter cassette. The percentage of EGFP^+^ SMCs represents SMCs with NF-κB activation in the absence or presence of TNF (10 ng/mL). The groups were compared by one-way ANOVA followed by Tukey’s post hoc analysis. P-value: n.s., not significant; *p<0.05; **p<0.01; ***p<0.001; ****p<0.0001.

We next examined NF-κB activation by assessing p65 nuclear translocation via immunostaining. Approximately 5% of SMCs exhibited basal nuclear p65 localization under static culture conditions. TNF treatment increased this to ∼25%, but translocation remained unchanged between static and stretched cells with or without TNF **(Figure 2D)**. Since the proportion of p65-positive nuclei was unchanged, this indicates that physiological stretch does not act by blocking p65 nuclear entry. Because *IKBKB* was nonetheless required for the stretch-induced suppression of *CCL2*, we suggest that physiological stretch constrains NF-κB output at a step downstream of translocation.

To obtain an independent, functional readout of NF-κB activation, we generated SMCs stably expressing an integrated lentiviral NF-κB-EGFP reporter (three κB elements driving a destabilized EGFP), which integrates NF-κB transcriptional output over time in single living cells rather than scoring localization at a fixed timepoint **(Supplementary Figure 1)**. EGFP-positive cells rose from 11% under basal conditions to 34% after TNF, with a shift toward higher per-cell intensity **(Figure 2E)**. Both fractions exceeded the corresponding immunostaining values (∼5% and ∼25%), as expected for an integrated transcriptional readout. Together, these results indicate that physiological stretch suppresses NF-κB-dependent inflammatory gene expression in SMCs, while a subset of cells exhibits basal or TNF-inducible NF-κB activation.

### 3.3 Transcriptional changes in proliferative and contractile programs precede detectable functional change

To determine whether stretch-induced cell cycle gene expression translated into functional changes, MKI67 immunostaining and EdU incorporation were assessed. MKI67-positive SMCs were reduced after 15% stretch compared with 10% stretch, with no difference between 10% stretch and static conditions **(Supplementary Figure 2A)**; EdU incorporation was unchanged at either stretch intensity, including after prolonged (24 h) 10% stretch **(Supplementary Figure 2B-C)**. Similarly, the modest increase in contractile gene expression under physiological stretch did not translate into altered contractile function as collagen gel contraction over 48 h was comparable between 10% stretched and static SMCs. Treatment with the positive control, TGFβ1, increased collagen gel contraction in all conditions to a similar extent, indicating that prior stretch did not alter the contractile response to TGFβ1 **(Supplementary Figure 2D-E)**. Together, stretch-induced transcriptional changes in proliferative and contractile programs were not accompanied by corresponding functional effects under our experimental conditions.

### 3.4 Single-cell RNA sequencing identifies distinct SMC subpopulations with a stable architecture across culture conditions

The presence of a constitutively NF-κB-active SMC subset at baseline, and its expansion under TNF, indicated that pathway activation is heterogeneous across cultured cells. To resolve this heterogeneity and determine whether the anti-inflammatory effect of physiological stretch is uniform across the SMC population or restricted to particular subsets, we performed scRNA-seq of SMCs cultured under static, 10% stretch, or TNF-stimulated conditions. Nine transcriptionally distinct clusters (C0–C8) were identified **(Figure 3A and Supplementary Table 4)**. Cluster architecture and distribution of cells was similar across static, stretch, and TNF conditions **(Supplementary Table 5)**, indicating that mechanical and inflammatory stimuli modulate transcriptional states within existing SMC subpopulations rather than by redistributing cells between states.

**Figure 3:**
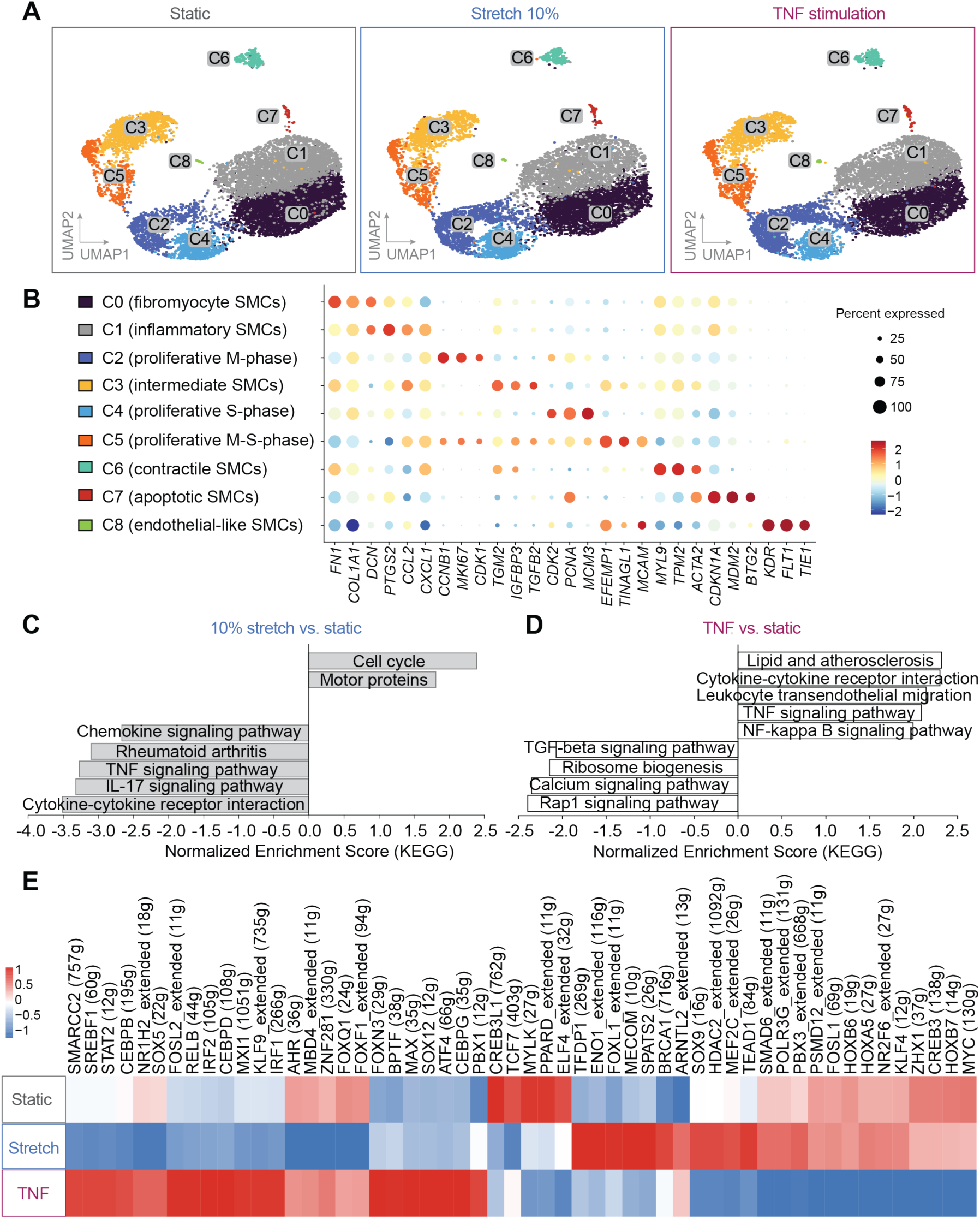
Transcriptomic characterization of mechanical stretch and TNF in SMCs by scRNA-seq. **A.** Outline of the experiment. Human SMCs were exposed to static conditions (24 h), mechanical stretch (10%, 6 h) or TNF stimulation (10 ng/mL, 24 h). Uniform Manifold Approximation and Projection (UMAP) visualization of the phenotypic diversity of SMCs after static culture, stretch and TNF stimulation. Each dot represents a single cell. Nine cell clusters were identified in human SMCs (0.3 resolution) as follows: C0 (fibromyocyte SMCs), C1 (inflammatory SMCs), C2 (proliferative M-phase), C3 (intermediate SMCs), C4 (proliferative S-phase), C5 (proliferative M-S-phase), C6 (contractile SMCs), C7 (apoptotic SMCs), C8 (endothelial-like SMCs). **B.** Dot plot showing the expression of three genes defining each cell cluster shown in A. Color scale indicates log2 expression, and the size represents the percentage of cells. Enrichment analysis of differentially expressed genes (DEGs) between **C.** 10% stretch and static conditions or **D.** TNF and static by KEGG. **E.** Heatmap of regulon activity for the three conditions from SCENIC gene regulatory network inference. Color intensity indicates z-scaled area under the curve (AUC). g indicates the number of genes on each regulon.

Cluster identity was assigned based on established marker genes **(Figure 3B)**. C0 (fibromyocyte SMCs) expressed extracellular matrix (*FN1, COL1A1, DCN*) and pro-inflammatory genes; C1 (inflammatory SMCs) showed high expression of inflammatory genes (*PTGS2* and *CCL2*); C2 (proliferative, M-phase) and C4 (proliferative, S-phase) were distinguished by cell cycle markers (*MKI67* and *CDK1*; *CDK2, PCNA*, and *MCM3*, respectively), C5 represented a transitional M-S-phase proliferative state; C3 (intermediate SMCs) shared the high inflammatory gene expression characteristics of C1, but also displayed higher expression of cell adhesion and migration-associated genes (e.g., *TGM2, IGFBP3, TGFB2*); C6 (contractile SMCs) expressed contractile genes (*MYL9, TPM2, ACTA2*). Clusters C7 and C8 were very small (≤1% of all cells). C7 (apoptotic SMCs) expressed *CDKN1A, MDM2*, and *BTG2*; and C8 (endothelial-like SMCs) expressed the endothelial markers *KDR, FLT1*, and *TIE1*. Representative gene markers for each cell cluster are shown in the **Supplementary Figure 3.** This annotation indicates that cultured human SMCs comprise a heterogeneous mixture of fibromyocyte, inflammatory, proliferative, and to a lesser extent, contractile states.

### 3.5 Stretch and TNF drive opposing transcriptional and regulatory programs across SMC subpopulations

Having established a stable population structure across conditions, we next examined the pathways and transcriptional regulators underlying the condition-specific expression changes within it. KEGG pathway enrichment analysis was performed on the differential expression analysis of 10% stretch versus static and of TNF versus static comparisons across all cells **(Figure 3C-D, and Supplementary Tables 6-7)**. Physiological stretch upregulated cell cycle-associated pathways and downregulated multiple inflammatory pathways, including chemokine, TNF, and IL-17 signaling, and cytokine-cytokine receptor interaction **(Figure 3C)**. TNF stimulation, as expected, upregulated pathways directly relevant to vascular inflammation, including such categories as lipid and atherosclerosis, cytokine-cytokine receptor interaction, leukocyte transendothelial migration, TNF and NF-κB signaling, while downregulating TGFβ signaling, ribosome biogenesis, calcium and Rap1 signaling **(Figure 3D)**. Together, these analyses indicate that physiological stretch and TNF drive largely opposing transcriptional programs. To identify the transcription factors underlying these condition-specific programs, we inferred regulon activity using SCENIC **(Figure 3E)**. Inflammatory regulators, including the NF-κB family member RELB together with CEBPB, CEBPD, IRF1, IRF2, and STAT2, showed their highest activity under TNF stimulation, supporting TNF as the principal driver of inflammatory transcription in this dataset. Notably, the activity of these regulons was lowest under physiological stretch, falling below the static baseline rather than merely below the TNF-stimulated state, indicating that physiological stretch reduces inflammatory transcription-factor activity rather than leaving it unchanged. By contrast, physiological stretch selectively increased the activity of a distinct group of regulons including MEF2C, TEAD1, SMAD6, and HDAC2, transcription factors linked to smooth muscle lineage maintenance, mechanotransduction, and chromatin regulation. Additional condition-associated regulons are provided in **Supplementary Table 8**. This result is in line with the working model that physiological stretch constrains NF-κB transcriptional output rather than its nuclear entry, and identifies the inflammatory regulons suppressed under stretch as candidate mediators of this effect.

### 3.6 Physiological stretch suppresses TNF-induced inflammatory genes across major SMC subpopulations

To determine whether the anti-inflammatory effect of stretch is uniform across the SMC population or restricted to a subset of cells, we compared differentially expressed genes (DEGs) from the 10% stretch versus static and TNF versus static comparisons within each cluster (**Figure 4A-F**, clusters C0-C5). Plotting these two comparisons against one another resolves genes that are induced by TNF but suppressed by physiological stretch (lower-right quadrant). Across clusters C0-C5, multiple TNF-induced inflammatory genes, including *IL1B, CCL2, CXCL1, CXCL8, IL6*, and *CSF3*, fell within this quadrant, indicating suppression by physiological stretch. Clusters C6–C8 contained too few cells for reliable differential expression analysis and were not assessed **(Supplementary Table 9)**.

**Figure 4.**
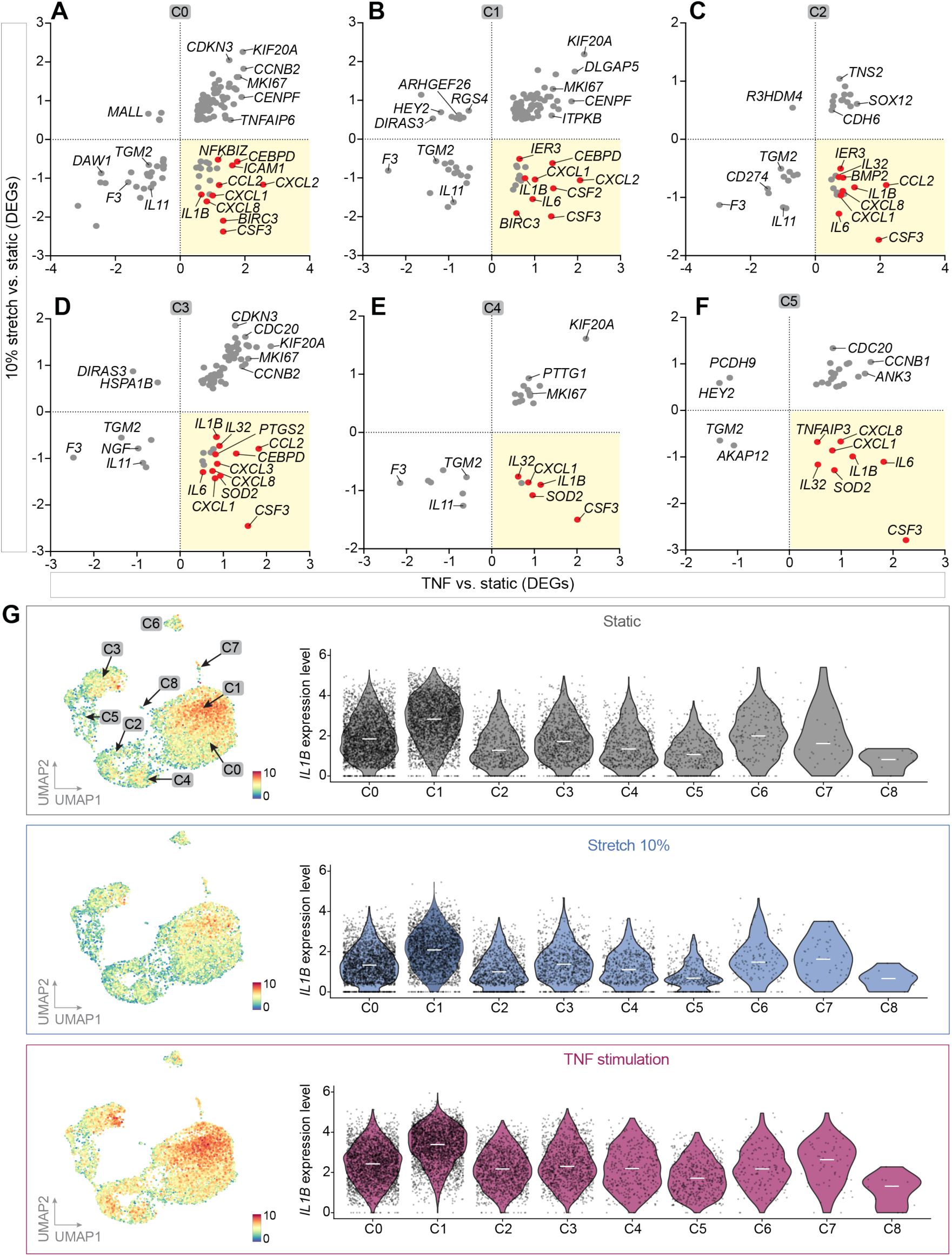
Physiological stretch attenuates pro-inflammatory signaling across clusters in cultured SMCs. **A-F** The log2 fold-change of shared differentially expressed genes (DEGs) with false discovery rate adjusted p-values (padj) <0.05 and log2 fold-change greater than 0.5 or less than −0.5 between stretch vs. static (*y*-axis) and TNF vs. static conditions (*x*-axis) in the different cell clusters C0 to C5 were plotted against each other. Static conditions (24 h), mechanical stretch (10%, 6 h) and TNF stimulation (10 ng/mL, 24 h) treatments were compared. Genes significantly downregulated by 10% stretch and upregulated by TNF are highlighted in the lower right (yellow) quadrant. Inflammatory gene markers are shown in red color. **G.** Normalized expression levels of *IL1B,* a pro-inflammatory gene (as example), in each cell cluster during static conditions, 10% mechanical stretch and TNF stimulation after scRNA-seq analysis.

In agreement with this signature, *IL1B* expression was reduced under physiological stretch relative to static and TNF-stimulated conditions across the major SMC clusters **(Figure 4G)**; the same effect was observed for other NF-κB target genes (*CCL2, IL6*; **Supplementary Figure 4**). Together, these findings indicate that physiological stretch suppresses pro-inflammatory gene expression across the major SMC subpopulations, counteracting TNF-driven activation of NF-κB target genes.

### 3.7 Cultured SMC states overlap with modulated smooth muscle phenotypes in human atherosclerotic plaques

To assess the relevance of these *in vitro* SMC phenotypes to human disease, we integrated our scRNA-seq dataset with publicly available single-cell transcriptomes from human coronary and carotid atherosclerotic plaques^40–42^. The integrated plaque dataset resolved the expected major populations, including a mesenchymal supercluster, endothelial cells, multiple immune populations, and a minor neuronal cluster; the plaque mesenchymal compartment was further subdivided into seven clusters (pC0–pC6), comprising contractile SMCs (pC0), pericyte-like SMCs (pC1), intermediate SMCs (pC2), fibromyocyte SMCs (pC3), osteochondrogenic SMCs (pC4), fibroblasts (pC5), and undefined SMCs (pC6) **(Figure 5A)**. Most cultured human SMCs mapped to the plaque mesenchymal supercluster, particularly to its modulated clusters (pC2, pC3, pC4, and pC6) **(Figure 5B)**. Two subsets of cultured SMCs extended beyond the mesenchymal supercluster: *in vitro* clusters C3 and C5 skewed towards the plaque endothelial cluster, consistent with their expression of cell adhesion molecules, whereas C0, C1, and C7 skewed towards the plaque monocyte cluster, reflecting their inflammatory (C0, C1) or apoptotic (C7) gene expression. The proliferating *in vitro* clusters (C2, C4, C5) localized near the proliferative immune cluster, most likely reflecting shared cell cycle gene expression rather than a common lineage. The contractile cluster C6 mapped across the contractile (pC0), osteochondrogenic (pC4), and undefined (pC6) plaque SMC populations. Together, these analyses indicate that cultured SMCs are comparably heterogeneous as plaque SMCs and occupy overlapping transcriptional states. While they mapped predominantly to plaque fibromyocytes, they also spanned endothelial-, immune-, inflammatory-, and proliferative-associated phenotypes, indicating that standard culture conditions maintain SMCs across a spectrum of modulated states that overlaps substantially with those found in human atherosclerotic plaques.

**Figure 5:**
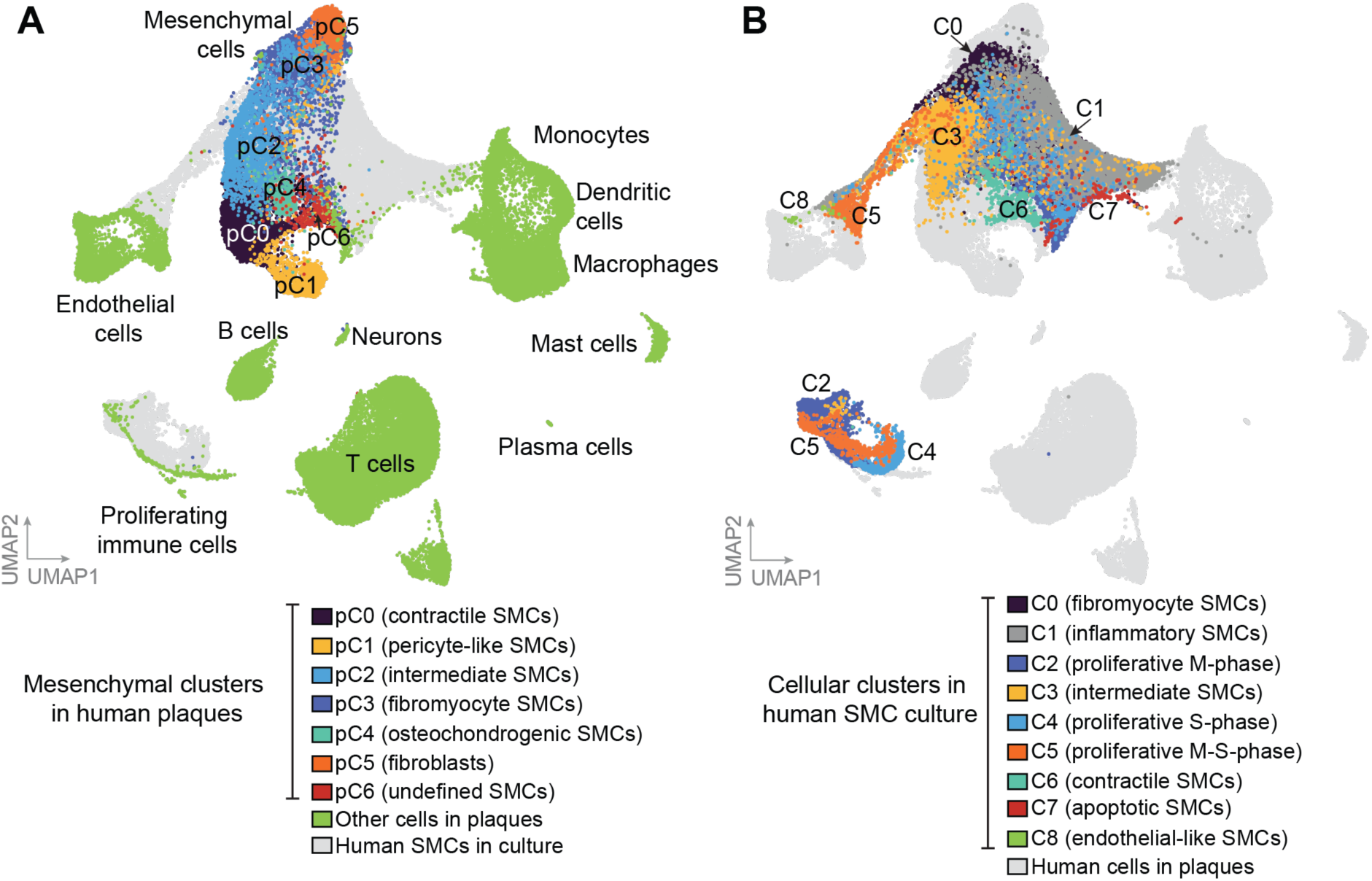
Integration of publicly available human atherosclerotic plaques scRNA-seq datasets with cultured human aortic SMCs from this study. **A.** Human plaque analysis revealed 7 mesenchymal cell clusters (pC0 to pC6 shown in the UMAP with different colors), where p stands for plaque. Analysis of human aortic SMCs in culture are shown in the background (light gray cluster). **B.** UMAP of cultured human aortic SMCs (C0 to C8) is shown in front with different colors and human atherosclerotic plaque cell populations are shown in the background (light gray clusters).

## 4. Discussion

Much of our mechanistic understanding of SMC biology derives from cells cultured under static conditions on rigid plastic. Although these models have been informative, they omit the cyclic mechanical forces that shape SMC behavior *in vivo*, and the phenotypic consequences of this omission have remained poorly defined^14,18^. Here we show that human aortic SMCs respond to stretch through coordinated regulation of cell cycle, contractile and inflammatory gene programs, and that the intensity of stretch determines the direction of the response. Physiological stretch (10%) upregulated cell cycle genes, relative to static culture. However, these proliferative transcriptional changes were not mirrored functionally: neither EdU incorporation nor was MKI67 positivity increased under physiological stretch. Likewise, the modest induction of contractile genes after physiological stretch did not enhance SMC contractility in collagen gel assays. This uncoupling of transcript from function is in agreement with previous reports^45,48–50^ and most likely reflects regulatory mechanisms in response to mechanical stimuli during the initial 6 h^51,52^ and post-transcriptional regulation^45,53,54^; sustained or repeated stretch may be required before these programs yield measurable functional effects.

The most consistent effect of physiological stretch was suppression of inflammatory gene expression. Physiological stretch attenuated both basal and TNF-induced expression of NF-κB target genes (e.g. *CCL2, ICAM1*) and lowered *IKBKB* transcript levels, whereas pathological stretch failed to attenuate inflammatory gene expression. Knockdown of *IKBKB*, which encodes the canonical NF-κB kinase IKKβ^55^, reduced baseline inflammatory gene expression and abolished the additional suppression of *CCL2* by stretch, indicating that IKKβ is required for this effect and that the two act in the same pathway. These findings align with a central role for NF-κB in SMC inflammation during atherogenesis^21,56–58^.

How stretch limits NF-κB output is less clear. The proportion of cells with nuclear p65 was unchanged by stretch, with or without TNF, so stretch does not act primarily by preventing p65 nuclear entry. This is counterintuitive with the fall in *IKBKB*, since IKKβ promotes IκB degradation and thus translocation^55^. A parsimonious interpretation is that IKKβ also acts beyond nuclear entry: IKKβ and related kinases phosphorylate p65 (e.g. Ser536) and other substrates governing transcriptional potency, so a partial reduction could blunt output while leaving bulk translocation intact^59,60^. These are, however, single-timepoint categorical measures that may miss graded dynamic changes in nuclear p65^61^, and the proposed post-translocation mechanism requires direct testing, for example by quantifying p65 phosphorylation and promoter occupancy. SCENIC regulon analysis supports this interpretation. Inflammatory regulons (RELB, CEBPB, CEBPD, IRF1, IRF2, STAT2) were most active under TNF and less active under physiological stretch, whereas stretch selectively increased regulons linked to smooth muscle lineage maintenance, mechanotransduction and chromatin regulation (MEF2C, TEAD1, SMAD6, HDAC2). Members of the MEF2 family have previously been implicated in stretch-induced transcriptional responses of vascular SMCs^62^, indicating a broader role for this family in SMC mechanical responses. MEF2 family may constrain NF-κB output indirectly, for example through competition for shared co-activators or reinforcement of non-inflammatory enhancers^63^.

In our hands 10% stretch suppressed human aortic SMC inflammatory response, whereas this suppression was lost at 15% stretch. This intensity dependence accommodates the apparently divergent literature^46,64^. Reports indicating that mechanical stretch activates NF-κB in vascular SMCs have generally used higher amplitudes, different waveforms, or stiffer substrates^65–69^, conditions we would classify as pathological, whereas studies applying near-physiological stretch describe stable, non-inflammatory transcriptional programs^52–54^, in line with our findings. Two recent studies in comparable models support this interpretation. In primary human aortic SMCs, 9% cyclic stretch at 1 Hz was shown to sustain a stable ECM- and TGFβ-related transcriptional program over 4-72 h, whereas unstretched controls exhibited progressive drift^52^, consistent with the notion that static culture represents a modulated baseline which physiological stretch attenuates. Conversely, low-amplitude high-frequency stretch of human intestinal SMCs, classified as pathological on the basis of frequency rather than amplitude, induced *CCL2, IL-6* and additional inflammatory mediators within 6 h^67^. In our aortic SMCs, 15% stretch did not reproduce the suppression seen at 10%, consistent with a loss of the physiological set-point at higher stretch. In both cases pathological is defined by failure to maintain the physiological, low-inflammatory set-point, not by active induction. In line with the basal subset of p65-positive cells and their heterogeneous TNF responses, scRNA-seq resolved nine transcriptional states with an extremely low proportion of contractile cells. Most cultured SMCs mapped to fibromyocytes and other modulated phenotypes of human atherosclerotic plaques, supporting the disease relevance of the model. Cluster architecture was preserved across conditions, indicating that stretch and TNF modulate SMC states within existing subpopulations. Physiological stretch suppressed inflammatory genes across the major clusters (C0–C5), counteracting TNF-driven induction and indicating that the anti-inflammatory effect is distributed broadly across these subpopulations rather than confined to a single state.

Together, these results indicate that static culture favors an intrinsically modulated, pro-inflammatory SMC state overlapping transcriptionally with plaque phenotypes, and that physiological stretch attenuates this program. The majority of cultured SMCs map transcriptionally to modulated plaque phenotypes rather than to contractile medial SMCs. A key implication of this work is that standard static culture should not be treated as a healthy control for vascular SMCs. Because the static baseline is transcriptionally modulated, inflammation-prone, and overlaps with plaque SMC states, it is more appropriately regarded as a disease-like model than as a physiological reference, and physiological mechanical loading is required to move SMCs toward a less-inflammatory phenotype. Incorporating physiological mechanical forces may therefore yield *in vitro* models that better represent healthy vasculature and are more predictive of SMC behavior in disease.

## 5. Limitations

Our data derive from a single donor SMC line, in line with recent single-donor aortic^52^ and visceral^67^ SMC studies; multi-donor, mixed-sex replication will be required to determine how age, sex, and cardiovascular risk background modulate the response. Stretch was applied for a single 6 h interval; longer and directional protocols may yield different outcomes^52^. NF-κB target regulation was assessed largely at the mRNA level, and the proposed post-translocation mechanism awaits protein-level confirmation; the NF-κB-EGFP reporter could not be applied under stretch, because lentiviral transduction was incompatible with the strain device. Future work should define how stretch shapes NF-κB activity, including p65 phosphorylation and translocation dynamics, IKKβ activity, and the upstream mechanotransduction pathways that initiate the response.

## 6. Conclusion

Standard static culture should not be regarded as a physiological reference for vascular SMCs but as a disease-like state that overlaps transcriptionally with modulated plaque populations. Physiological cyclic stretch attenuates this state and constrains inflammatory transcriptional output, including that of NF-κB, at a step downstream of p65 nuclear entry. Incorporating physiological mechanical loading into *in vitro* SMC models is therefore likely to yield systems more representative of the healthy vessel wall and more informative for atherosclerosis research.

## CRediT authorship contribution statement

Lise F. Jensen: Conceptualization, Investigation, Formal analysis, Visualization, Writing-original draft, Writing-Review & Editing. Anton Markov: Formal analysis, Visualization, Writing-Review & Editing. Laura Alonso-Herranz: Investigation, Formal analysis, Visualization, Writing-Review & Editing. Diana Sharysh: Formal analysis, Visualization, Writing-Review & Editing. Emil A. Thomsen: Supervision, Formal analysis, Writing-Review & Editing. Jacob G. Mikkelsen: Supervision, Writing-Review & Editing. Jacob F. Bentzon: Conceptualization, Formal analysis, Supervision, Funding acquisition, Writing-original draft, Writing-Review & Editing. Julián Albarrán-Juárez: Conceptualization, Investigation, Formal analysis, Supervision, Visualization, Funding acquisition, Writing-original draft, Writing-Review & Editing.

## Data availability statement

Raw and processed bulk and single-cell RNA sequencing data are available in the ArrayExpress collection of BioStudies repository (http://www.ebi.ac.uk/biostudies) under accession numbers E-MTAB-15611 and E-MTAB-15614, respectively. Any additional information required to reanalyze the data reported in this study, including the code, is available from the corresponding author upon reasonable request.

## Acknowledgments

We thank Lisa Maria Røge and Dorte Qualmann for their excellent technical assistance. Flow cytometry was performed at the FACS Core Facility, Aarhus University, Denmark. We are grateful to Martin Schwartz and Brian Coon (Yale University, USA) for generously providing the NF-κB reporter vector. We thank Miguel Angel del Pozo and Laura Sotodosos-Alonso (CNIC, Spain) for valuable discussions and insights into mechanotransduction. We also thank the BD Biosciences team, particularly Soudabeh Rad Pour and Jouni Rantanen, for training and insightful discussions on the Rhapsody single-cell RNA sequencing system. Finally, we acknowledge GenomeDK and Aarhus University for providing the computational resources and support essential to this research.

---

**Supplementary Figure 1:**
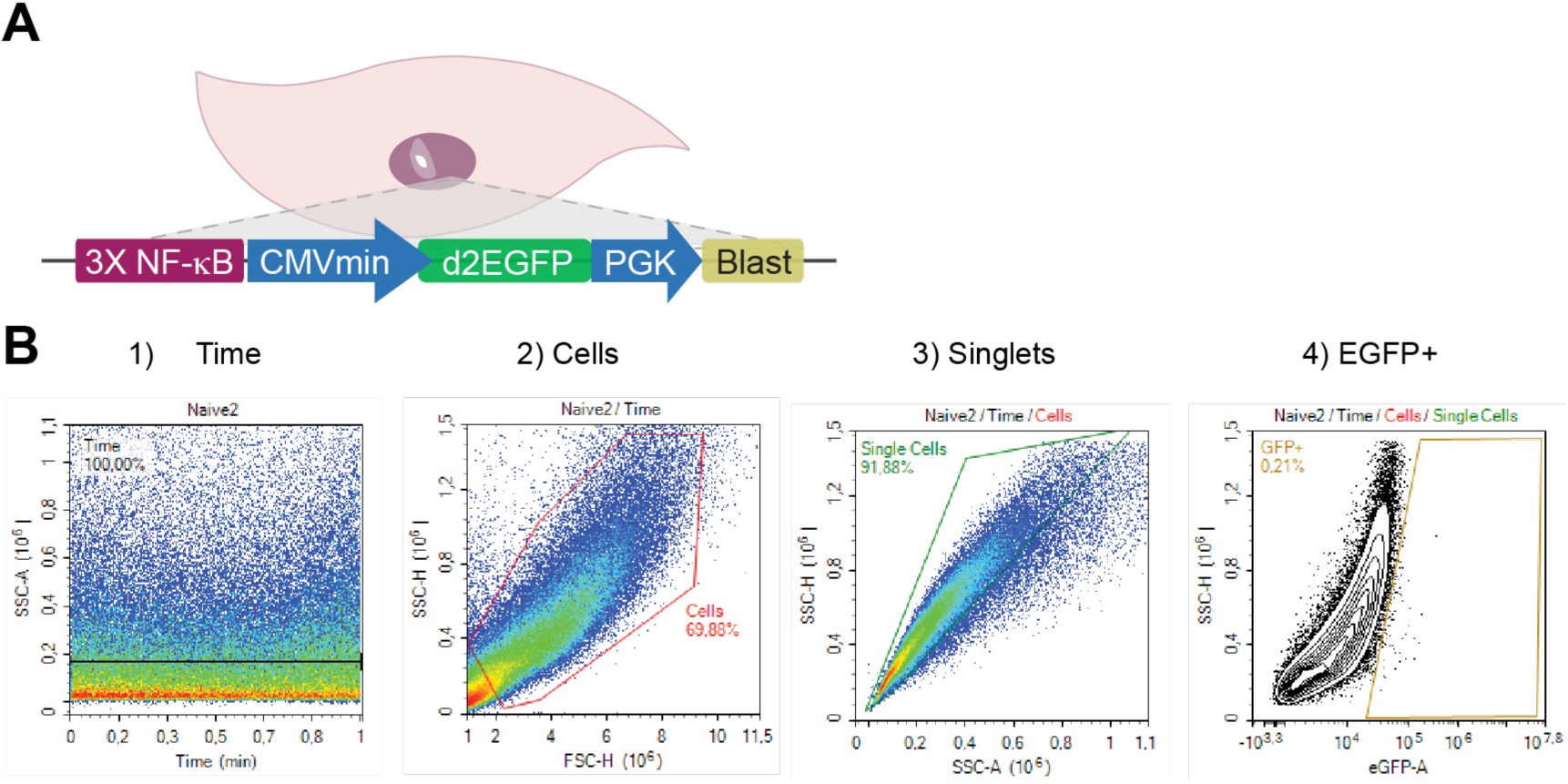
NF-κB-EGFP reporter assay. **A.** Schematic of the NF-κB reporter, where three NF-κB-response elements control the activation of a CMV minimal (CMV_min_) promoter controlling the expression of a destabilized EGFP gene (d2EGFP), and a phosphoglycerate kinase (PGK) promoter constitutively expressing a blasticidin resistance gene (Blast). Lentiviral transduction was used to deliver the reporter cassette in human SMCs. **B.** Representative gating strategy in flow cytometry using non-transduced (Naïve) SMCs to define the EGFP^+^ SMCs. 1) Time versus SSC-A was plotted to ensure stable flow. 2) SMCs were gated on SSC-H x FSC-H to exclude debris. 3) Singlets were identified by SSC-H x SSC-A. 4) EGFP^+^ SMCs were gated on SSC-H x EGFP. SSC: side scatter, FSC: forward scatter, A: area, H: height.

**Supplementary Figure 2:**
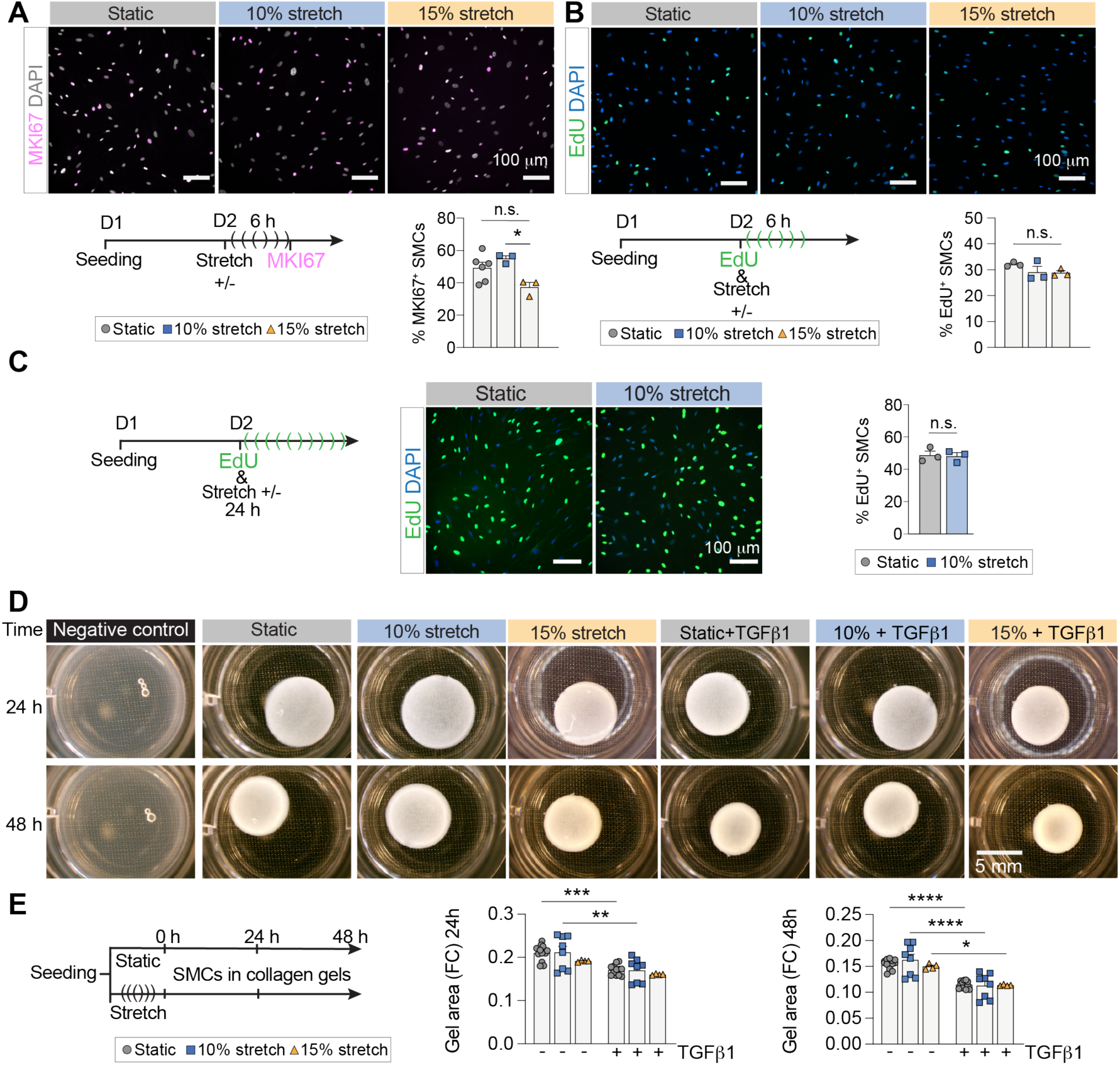
Mechanical stretch does not alter proliferation or contractile phenotype of SMCs cultured on collagen I. **A.** Representative images of SMCs immunostained with MKI67-Alexa Fluor 488 (magenta) and DAPI (gray). Cells were fixed immediately after being exposed to different stretch conditions and stained. The bar graph shows the percentage of MKI67-positive SMCs relative to the total nuclei count (*n*=3-6 independent replicates per group). **B.** Representative images of SMCs labeled with EdU-Alexa Fluor 488 (green) and DAPI (blue). Cells were cultured for 24 h, then EdU was added to the culture, and cells were subsequently exposed to the different stretch conditions (as indicated) for 6 h or left under static conditions. After stretching, the cells were fixed and EdU-labeling was detected. The bar graph shows the percentage of EdU-positive SMCs relative to the total nuclei count (*n*=3 independent replicates per group). **C.** EdU was added to the culture and SMCs were subsequently exposed to 10% stretch or left under static conditions for 24 h. After stretching, the cells were fixed and EdU-labeling detected. Shown are representative images of SMCs labeled with EdU-Alexa Fluor 488 (green) and DAPI (blue) and the percentage of EdU-positive SMCs relative to the total nuclei count (*n*=3 independent replicates per group). The groups were compared by unpaired t-test, P-value: n.s., not significant. **D.** Shown are representative images of collagen-I gels, without cells (negative control), or with SMCs subjected to the different experimental conditions (at least *n*=4 independent replicates per group). **E.** Schematic of the experimental design for the contractility assay. Cells were left static or subjected to 10% or 15% stretch (6 h) and embedded into collagen-I gels. Cells were either non- or TGFβ1-stimulated. The quantification of the gel area is shown as the FC compared to the negative control (*n*=8 independent replicates per group). The groups were compared by one-way ANOVA followed by Tukey’s *post hoc* analysis, adjusted p-value: n.s., not significant; *p<0.05; **p<0.01. D: Days. Scale bars:100 µm (for A-B), 5 mm (for D).

**Supplementary Figure 3:**
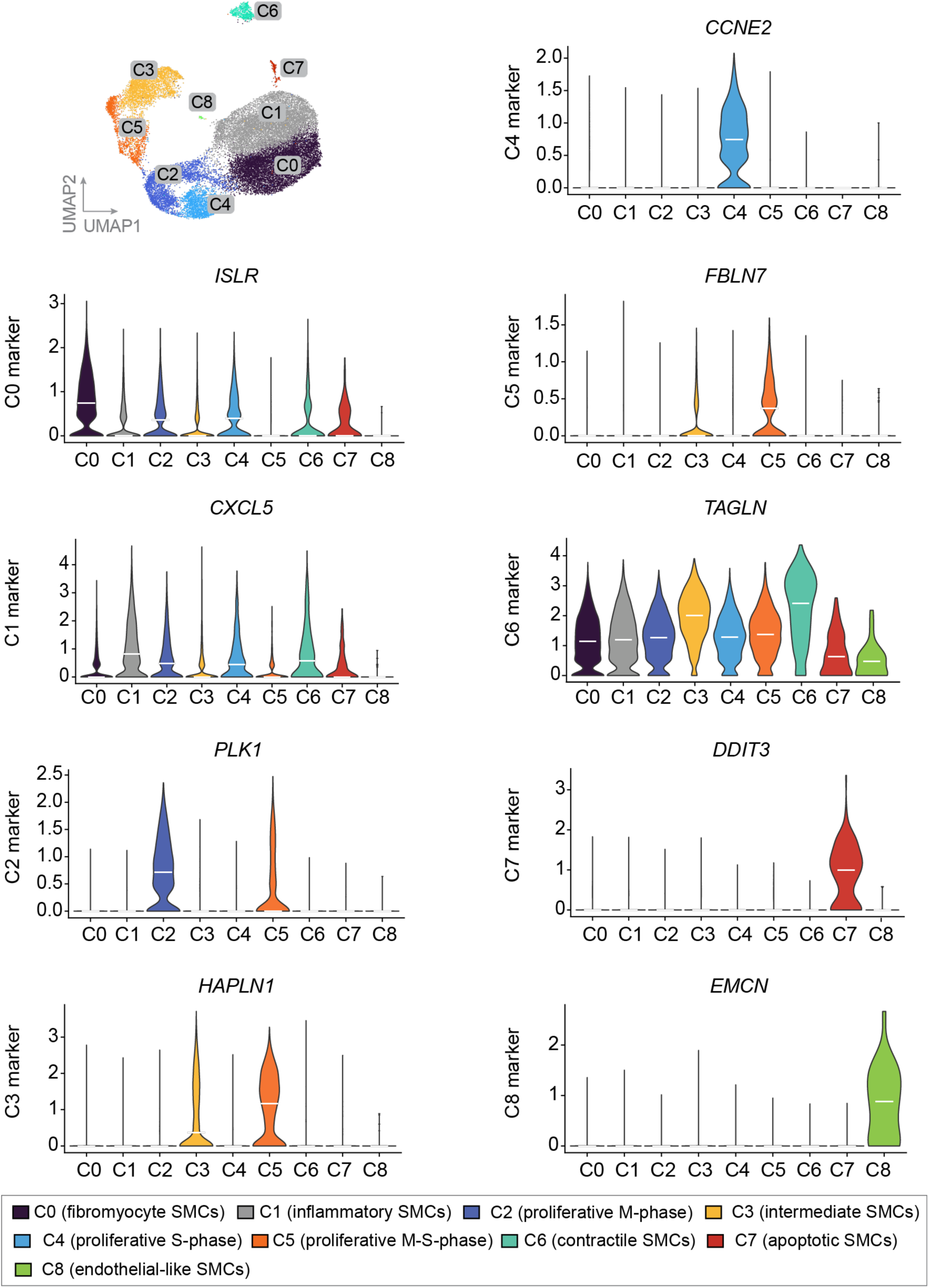
Examples of marker genes for the different cell clusters. Violin plots show the distribution of normalized expression for the selected marker genes for each cluster obtained in the scRNA-seq analysis. Expression is shown as the mean of normalized expression of these genes in each cluster. Genes with a false discovery rate adjusted p-values (padj) <0.05 and log2 fold-change greater than 0.25 or less than −0.25.

**Supplementary Figure 4:**
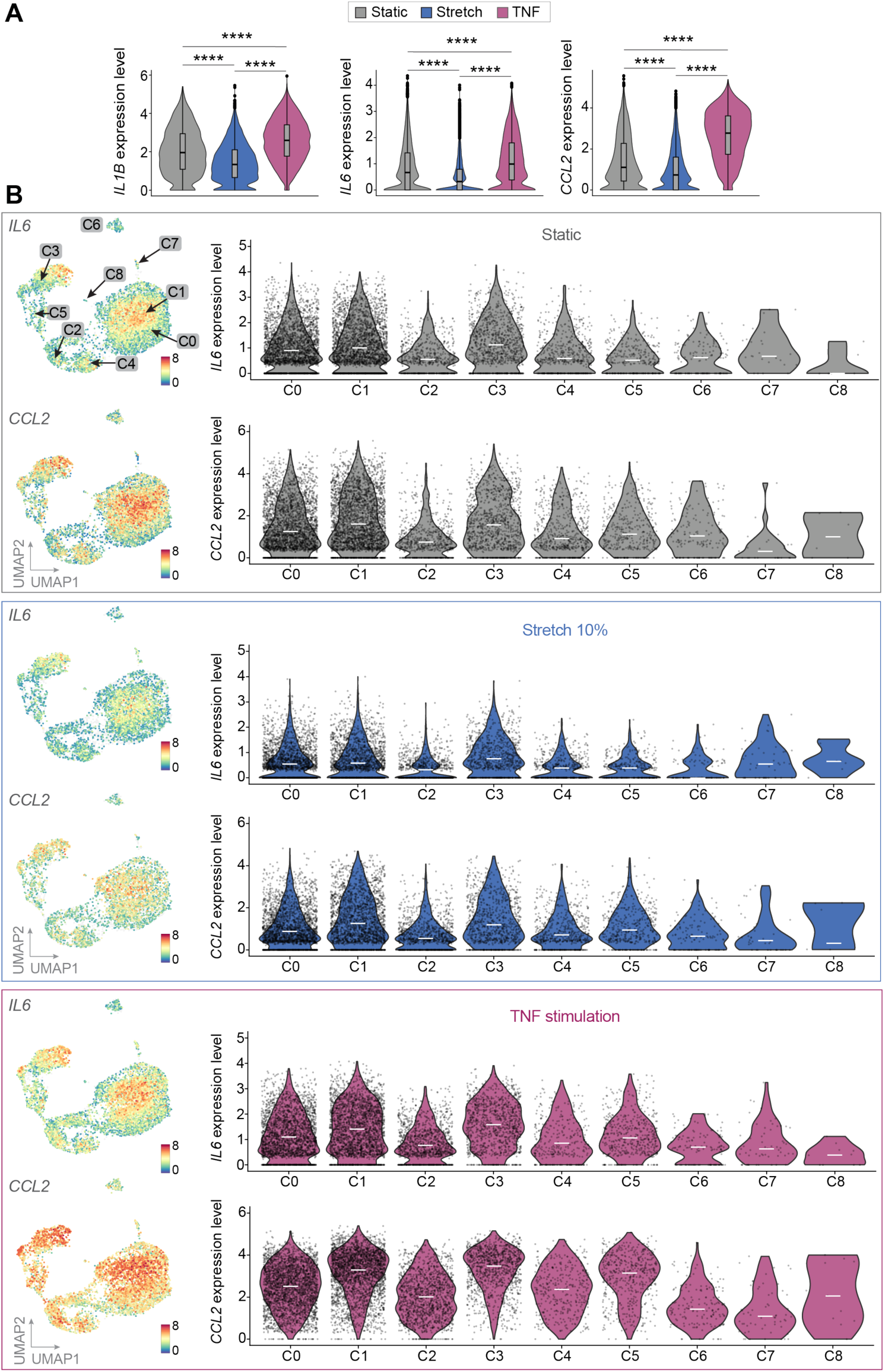
Expression of selected pro-inflammatory genes in scRNA-seq analysis. **A.** Mean of normalized expression across all cells of selected genes (as indicated) for each condition. Groups were compared by the Wilcoxon’s test. P-value: ****p<0.0001. **B.** Single-cell expression (UMAP visualization) of selected proinflammatory markers genes (*IL6,* and *CCL2)*, and normalized expression across SMCs clusters under static conditions (24 h), mechanical stretch (10%, 6 h) and TNF stimulation (10 ng/mL, 24 h). Color scale indicates log2 expression. C0 (fibromyocyte SMCs), C1 (inflammatory SMCs), C2 (proliferative M-phase), C3 (intermediate SMCs), C4 (proliferative S-phase), C5 (proliferative M-S-phase), C6 (contractile SMCs), C7 (apoptotic SMCs), C8 (endothelial-like SMCs).

**Supplementary Figure 5:**
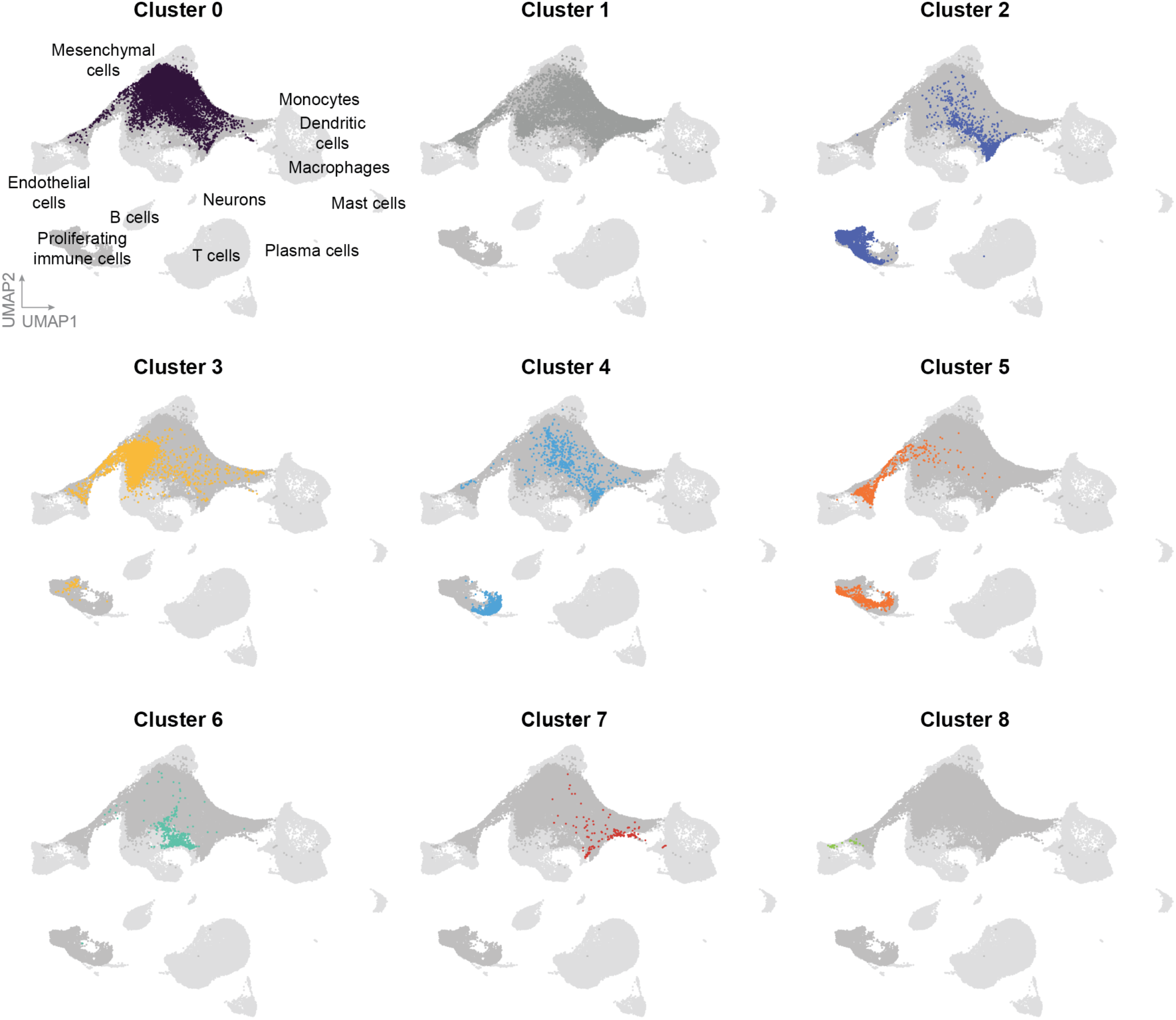
Individual clusters (C0-C8) from cultured SMC scRNA-seq and their integration with human plaque scRNA-seq. Shown are UMAP plots for each cell cluster identified in human aortic SMCs (C0-C8, different colors). C0 (fibromyocyte SMCs), C1 (inflammatory SMCs), C2 (proliferative M-phase), C3 (intermediate SMCs), C4 (proliferative S-phase), C5 (proliferative M-S-phase), C6 (contractile SMCs), C7 (apoptotic SMCs), C8 (endothelial-like SMCs). Integration of human coronary and carotid atherosclerotic plaques scRNA-seq datasets are shown in the background in a light gray color.

